# Whole plastid genome-based phylogenomics supports an inner placement of the *O. insectifera* group rather than a basal position in the rapidly diversifying *Ophrys* genus (Orchidaceae)

**DOI:** 10.1101/2020.12.16.423003

**Authors:** Joris A. M. Bertrand, Anaïs Gibert, Christel Llauro, Olivier Panaud

**Author notes:** Joris A. M. Bertrand, Laboratoire Génome & Développement des Plantes (UMR 5096 UPVD/CNRS), Université de Perpignan Via Domitia, Bâtiment T, 58 avenue Paul Alduy, 66860 Perpignan Cedex 09, France. ***Joris Bertrand*** conducted fieldwork, analyzed the data and wrote the manuscript. ***Anaïs Gibert*** contributed to fieldwork and manuscript writing. ***Christel Llauro*** generated the long-read data set. ***Olivier Panaud*** contributed to research funding and manuscript writing.

## Abstract

Some lineages of the Orchid genus *Ophrys* exhibit among the highest diversification rates reported so far. As a consequence of a such intense and rapid evolution, the systematics and the taxonomy of this genus remains unclear. A hybrid assembly approach based-on long- and short-read genomic data allowed us to outperform classical methods to successfully assemble whole plastid genomes for two new *Ophrys species*: *O*. *aymoninii* and *O*. *lutea*. Along with three other previously *Ophrys* plastid genome sequences, we then reconstructed the first whole plastome-based molecular phylogeny including representatives of the three mains recognized *Ophrys* lineages. Our results support the placement of the *O*. *insectifera* clade as sister group of ‘non-basal *Ophrys*’ rather than a basal position. Our findings corroborate recent results obtained from genomic data (RAD-seq and transcriptomes) but contrast with previous ones. These results therefore confirm that molecular phylogenetic hypotheses based on a limited number of *loci* (e.g. *nrITS, matK, rbcL*) may have provided a biased picture of phylogenetic relationships within *Ophrys* and possibly other plant taxa.

## 1. Introduction

Among the most speciose family of flowering plants that orchids (Orchidaceae) form, some lineages of the genus *Ophrys* display among the highest diversification rates ever reported (Givnish et al. 2015; Breitkopf et al. 2015). The adaptive radiation that *Ophrys* experience is likely to be due to their unusual pollination strategy (by sexual swindle) that leads to high levels of specialisation of these plants to their insect pollinators and favour evolutionary divergence (see Baguette et al. 2020 for a recent review). The systematic relationships of such fast and importantly diversifying groups are difficult to infer for two reasons. Firstly, because recent divergent times often renders molecular signal of lineage delineation undetectable or at least ambiguous (incomplete lineage sorting) and because emerging species are still particularly prone to introgressive hybridization and reticulate evolution.

The systematics and the taxonomy is particularly problematic in *Ophrys* for which different authors recognize a number of species ranging from 9 to 354 (see Bateman et al. 2018; Bateman 2018, Bertrand et al., 2021). In particular, contrasting results still make the phylogenetic position of the (three) main lineages (sometimes considered as subgenera) debated. Several molecular phylogenetic studies show that the genus *Ophrys* is basically subdivided in three main sub-lineages (sometimes considered as subgenera): a first clade formed by the *Ophrys insectifera* group (also defined as group A since the study of Devey et al. 2008), a second clade consisting of the groups B to E (*O. tenthredinifera* (B), *O. speculum* (C), *O. bombyliflora* (D), called ‘archaic *Euophrys*’ by Tyteca and Baguette (2017), plus the so-called *Pseudophrys* group (E) and a third clade to which belong the groups F to J (*O. apifera* (F), *O. sphegodes* (G), *O. fuciflora* (H), *O. scolopax* (I) and *O. umbilicata* (J), also called ‘recent *Euophrys*’). The terms *Euophrys* and *Pseudophrys* classify *Ophrys* according to the part of the pollinator insect’s body on which the pollinia are glued during pseudocopulation. *Pseudophrys* corresponds to *Ophrys* in which the pollinia are deposited on the abdominal region of the insect, while *Euophrys* corresponds to species in which the pollinia are deposited on its cephalic region (see Bertrand et al. 2021). As *Euophrys* is a taxonomically incorrect term to refer to *Ophrys* groups, ‘section *Ophrys*’ should be used as a contrast to section *Pseudophrys*. However, because the section *Ophrys* forms a paraphyletic group, we propose to use ‘basal *Ophrys*’ and *‘*non-basal *Ophrys*’ instead of ‘archaic *Euophrys’* and ‘recent *Euophrys*’ for clarity purpose.

Out of the studies that could not unambiguously resolve tree topology for the three main *Ophrys* lineages (e.g. Soliva et al. 2001; Tyteca and Baguette 2017) two contrasting hypothesis can be considered concerning the phylogenetic position of the *O. insectifera* (A) group. Most of the molecular phylogenetic hypotheses have (historically) rather supported a basal position (*T*_basal_) for the *O. insectifera* (A) group (Devey et al. 2008; Breitkopf et al. 2015, Zitoun et al. *in prep*). However, recent findings based on genomic data: SNPs derived from RAD-seq approaches (Bateman *et al*. 2018) or transcriptomes (Piñeiro Fernández et al. 2019) rather support that the group *insectifera* (A) is directly related to ‘non-basal *Ophrys*’ (groups F to H) both of which being sister to the clade comprising ‘basal *Ophrys*’ + *Pseudophrys* (groups B to E) (*T*_inner_).

In this study, we aim to reconstruct a phylogenomic hypothesis to test whether whole plastid genomic data rather support the inner placement of the *O. insectifera* (A) group or alternatively, its basal position in the *Ophrys* genus. So far, three *Ophrys* plastid genomes have been published: *O. iricolor* Desf. (or *O. fusca* subsp. *iricolor* (Desf.) K.Richt) and *O. sphegodes* Mill. (Roma *et al*., 2018) and *O. aveyronensis* (J.J.Wood) P.Delforge (or *O. sphegodes* subsp. *aveyronensis* J.J.Wood) (Bertrand *et al*., 2019), none of which are members of the *O. insectifera* (A) group. To fill this knowledge gap, we generated genomic data for *Ophrys aymoninii* (Breistr.) Buttler (or *O. insectifera* subsp. *aymoninii*, Breistr.) a representative of the *O. insectifera* clade, endemic to a spatially restricted geographic area in the South of the Massif Central (France). We also provide similar data for *O. lutea* Cav., 1793, a widespread Western Mediterranean *Pseudophrys* species.

We relied on a hybrid approach to assemble the whole plastid genomes of the two *Ophrys* taxa mentioned above. In brief, this consists in a combination of long reads (here, Oxford Nanopore Technologies reads that can span repeated DNA regions known to be difficult to assemble) with the low error rate of short (paired-end) reads (here, Illumina reads). Although relatively recent, such hybrid strategy was found to outperform classical approaches (as recently supported by Wang et al. 2018 and Scheunert et al. 2020). To do so, we used the Unicyler pipeline (Wick et al. 2017) which is able to analyse reads from both platforms simultaneously. Gene annotation and basic downstream analyses were then carried out as described in our former study (Bertrand et al. 2019, see also Appendix1).

## 2. Materials and Methods

### 2.1 Field sampling and sample processing

We collected fresh leaves from an individual of *O. aymoninii* and an individual of *O. lutea* near Causse-Begon, France (N 44.05252°; E 3.35898°) and Versols-Et-Lapeyre, France (N 43.898677°; E 2.933099°), respectively on 12-05-2018. As *O. aymoninii* is a nationally protected species in France, sampling was carried out under permit ‘Arrêté préfectoral n°2018-s-20’ issued by the ‘Direction Régionale de l’Environnement de L’Aménagement et du Logement (DREAL)’ from the ‘Région Occitanie’ on 11-06-2018. Back in the lab, samples were frozen and stored at −20°C until DNA extraction. We used a CTAB2X protocol to extract genomic DNA from the two specimens sampled (see Appendix 1 for details).

### 2.2 DNA sequencing, plastid genome reconstruction and gene annotation

#### 2.2.1 Long-read (Nanopore) sequencing

Four Nanopore sequencing libraries (two for each individual) were prepared with the SQK-LSK108 kit following the ONT “1D genomic DNA by ligation protocol” or the “1D gDNA long reads without BluePippin protocol” from 2730/3632 ng and 2964/2500 ng of unfragmented DNA for *O. aymoninii* and *O. lutea*, respectively (see Appendix 1 for detail). Long read sequencing was carried out from four FLO-MIN106D R9 flowcells (two for each individual) on a MinION (Oxford Nanopore Technologies, Oxford, UK) in the lab using MinKNOW v.1.15.1. FAST5 files were base-called with Albacore v.2.3.1 (see Appendix 1 for detail).

Adapters were removed with Porechop v.0.2.4 (https://github.com/rrwick/Porechop) with the –discard_middle option turned on. We then used Nanofilt v.2.7.1 (https://pypi.python.org/pypi/NanoFilt) to filter out reads shorter than 5 kb and bases with quality < 9 on both sides of reads. To facilitate assembly, we then extracted plastid reads by mapping short-reads onto a multi-fasta file comprising the three whole plastid genome sequences published for *Ophrys* species. As in Wang et al. (2018), we duplicated and concatenated each of the three sequences and included them in the reference set to avoid losing reads corresponding to the region where genomes were circularized. We then extracted reads that mapped onto this dataset with Minimap2 v.2.17 (Li, 2018).

#### 2.2.2 Short-read (Illumina) sequencing

Whole genomic libraries were prepared and sequenced in paired-end mode (2×150 bp, insert size: 350 bp) by Novogene Co., Ltd (HK) from 1.92 and 1.97 pg of DNA for *O. aymoninii* and *O. lutea*, respectively. Genomic DNA was extracted with the same protocol than the one used for long reads. Raw reads were trimmed with Trimmomatic v.039 (Bolger et al. 2014) and the resulting read quality was checked with FastQC v0.11.8 (Andrews et al. 2010). Plastid read extraction was carried out by mapping short-reads onto the *Ophrys* plastome dataset, as mentioned above, this time with bowtie2 v.2.3.4 (Langmead and Salzberg, 2012). As too high coverage is prone to disturb the assembly process, we subsampled the resulting read set to an expected coverage of 100X (*i*.*e*. by keeping 500,000 of both R1 and R2 reads assuming a plastid genome size of around 150 kb) before the assembly step.

#### 2.2.3 Plastid genome reconstruction and gene annotation

Hybrid *de novo* assembly was performed with both long- and short-reads simultaneously using default settings in Unicycler v0.4.9b (Wick et al. 2017). Gene annotation and alignments were performed as in our previous study (Bertrand et al. 2019). Some genes (*ndh*A to *ndh*K) exhibited significant differences in length and similarity, even between the closely related *Ophyrs* species considered here, probably as a result of pseudogenisation and were removed from the alignments for further analyses. In particular, *O. sphegodes, O. aveyronensis* and *O. aymoninii* presented truncation of most *ndh* genes and shared the loss of the partially duplicated gene of *ycf*1 and a truncation of *ndh*F gene as already reported by Roma et al. (2018), see also Appendix 4.

### 2.3 *Phylogenetic reconstruction*, concordance factors and tree topology tests

We used the Maximum Likelihood approach implemented in IQ-TREE v2.0.6 (Minh et al., 2020a) to reconstruct gene trees and species tree with 1,000 replicates (-B 1000) of Ultrafast Bootstrap Approximation (UFBoot) to assess nodes support. Species tree was constructed based on three data set: i) whole plastome, ii) genes and iii) CDS alignments. The genealogical concordance in the dataset was also quantified with gene concordance factor (gCF) and site concordance factor (sCF) (Minh et al., 2020b). In addition, tree topology tests implemented in IQ-TREE were performed to test whether the inferred species tree was rather consistent with the inner placement (*T*_inner_) or the basal (*T*_basal_) position of *O. aymoninii*. To further investigate which *loci* specifically supports one or the other topology and to what extent (by evaluating its phylogenetic signal), we then computed the difference in gene-wise Log-likelihood scores (ΔGLS, see Shen et al. 2017). For all phylogenetic analyses the plastid genome of another Orchidoideae species, *Platanthera japonica* (GenBank Accession no.: MG925368) was used as outgroup.

## 3. Characterization of the plastid genomes of *O. aymoninii* and *O. lutea* and comparison with previously published *Ophrys* plastid genomes

Following the approach described in our previous study (*i*.*e*. using short-reads (Illumina) and NOVOPlasty (Dierckxens et al. 2017, Bertrand et al. 2019)) we failed to infer a complete plastid genome sequence for *O. aymoninii*. For the two remaining species: *O. aveyronensis* and *O. lutea* we could obtain a single contig, only when providing a closely related reference sequence of *O. sphegodes* and *O. iricolor*, respectively (Table 1). The hybrid assembly implemented in Unicycler thus seems to outperform the short-read based approach as we obtained a single contig of expected length for all three species (without reference sequence). The two new *O. aymoninii* and *O. lutea* genomes were found to be very similar in size and structure as well as compared to the three previously published *Ophrys* plastome sequences. The plastid genomes of *O. aymoninii* and *O. lutea* are described in Appendix 2 and have been deposited on GenBank with accession numbers MW309825 and MW309826, respectively. Raw reads are also available from the European Nucleotide Archive (Study Primary Accession PRJEB42431/Secondary Accession ERP12689, see Appendix 1 for details).

**Table 1.** Comparative summary of the assembly of the plastid genomes of *Ophrys aveyronensis, O. aymoninii* and *O. lutea* based on the short-read approach (NOVOPlasty) and the hybrid approach (Unicycler).

|  | Short-read approach (NOVOPlasty) |  |  | Hybrid approach (Unicycler) |  | Reference |
| --- | --- | --- | --- | --- | --- | --- |
|  | Number of contigs (without reference) | Number of contigs (with reference) | Inferred sequence length (bp) | Number of contigs | Inferred sequence length (bp) |  |
| *Ophrys aveyronensis* | 3 | 1 | 146,816 | 1 | 146,816 | Bertrand et al. 2019 |
| *Ophrys aymoninii* | failed | 6-7 | NA | 1 | 146,674 | This study |
| *Ophrys lutea* | 3 | 1 | 150,338 | 1 | 150,284 | This study |

## 4. Phylogenetic relationships within the genus *Ophrys*

We found an overwhelming support for an inner placement (*T*_inner_) of the *O. insectifera* (A) group within the *Ophrys* genus. Whatever the alignment considered: whole-plastome, concatenation of gene/CDS *loci*, bootstrap values fully support (UFBoot = 100) a topology according to which the representative of the *O. insectifera* group: *O. aymoninii* is sister to the ‘non-basal *Ophrys*’ representatives: *O. sphegodes*/*O. aveyronensis*; the *Pseudophrys*: *O. lutea* and *O. iricolor* occupying a basal position (Figure 1). The gene and site concordant factor metrics (gCF and sCF) do not contradict bootstrap values even though they were found to be lower (especially gCF). This may be explained by very short branch lengths and the very limited amount of information contained in each gene/CDS sequence. All the tree topology tests also reject the topology consistent with the basal placement of *O. aymoninii* when compared to the one consistent with its inner placement (Table 2). The distribution of ΔGLS values (Figure 2 and Appendix 3) confirms that most of the genes also support this topology and when they do not, they only weakly support the alternative one.

**Figure 1.**
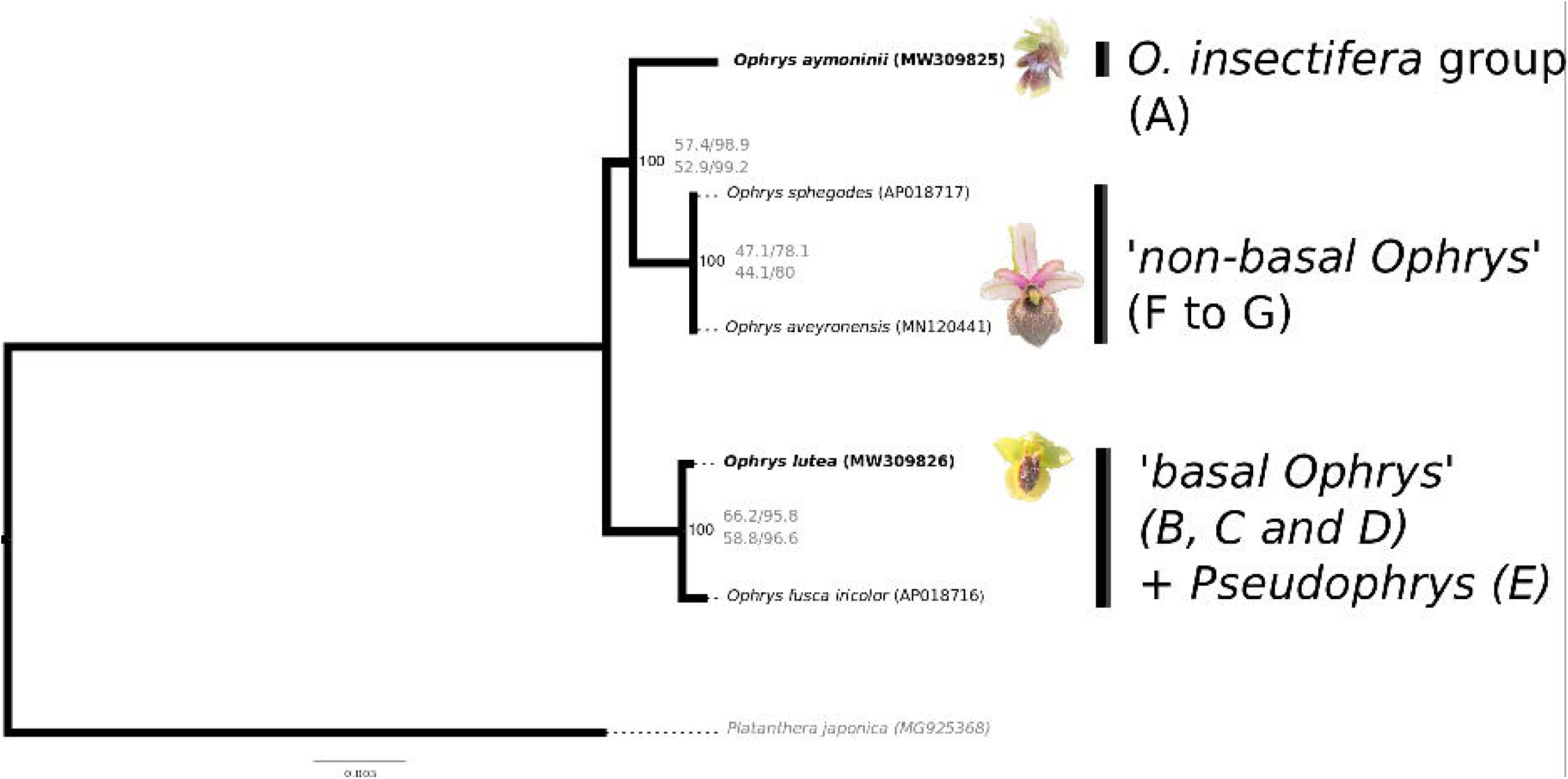
Maximum likelihood phylogenetic tree (as inferred with IQtree) from whole plastid genome sequence alignment. UFBoot (Ultrafast Bootstrap Approximation) values were 100 at each node whatever the alignment considered (whole plastome, genes, CDS). Values next to the bootstrap indicate gene Concordance Factor (gCF) and site Concordance factor (sCF) inferred from gene-(up) and CDS-based (down) ‘gene’ trees. *Platanthera japonica* (GenBank Accession no.: MG925368) is used as outgroup.

**Figure 2.**
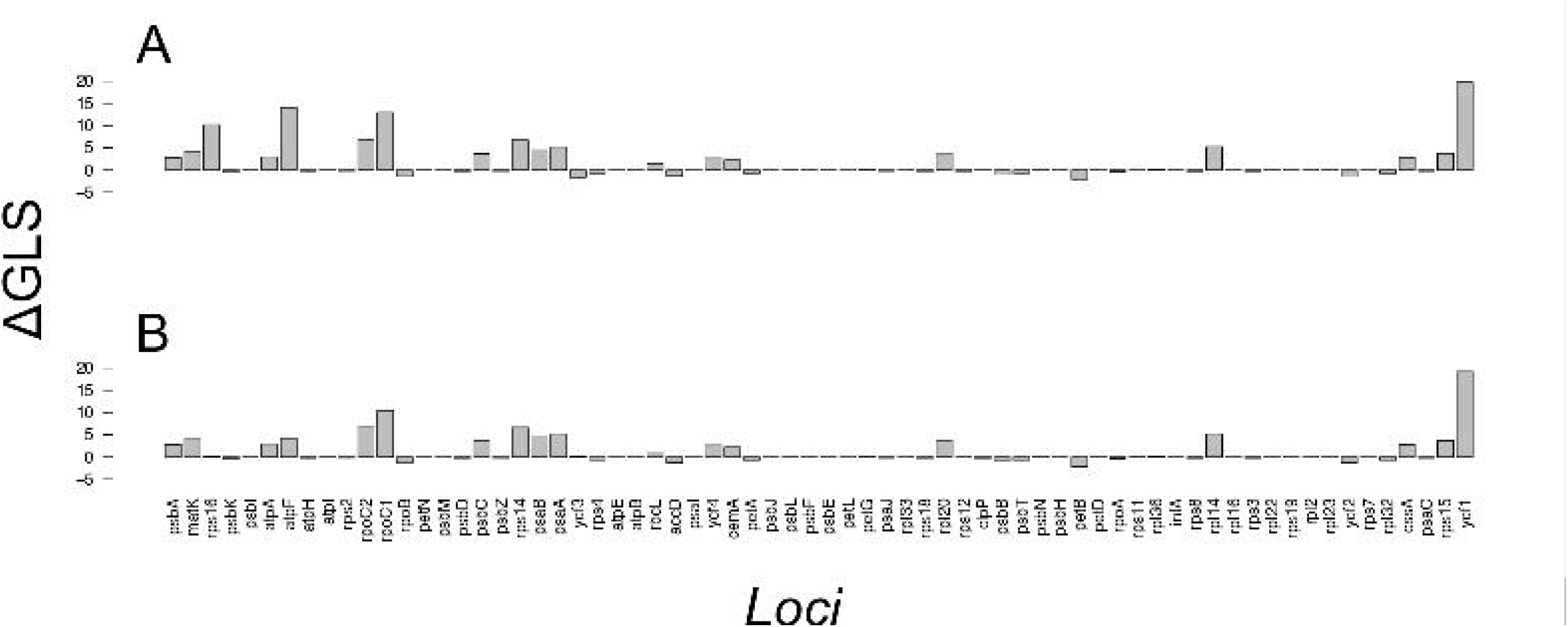
Genewise phylogenetic signal (ΔGLS) for *T*_inner_ versus *T*_basal_, the two alternative tree topologies for each gene (A) and CDS (B) along the plastid genome. Positive ΔGLS values support an inner placement of *O. aymoninii* whereas negative values rather support its basal placement.

**Table 2.** Summary of the tree topology test statistics performed to compare the inner placement hypothesis of the *O. insectifera* group (*T*_inner_) to the one supporting its basal position (*T*_basal_)

| Topology | logL | $\Delta L$ | bp-RELL | $p$-KH | $p$-SH | $p$-WKH | $p$-WSH | c-ELW | $p$-AU |
| --- | --- | --- | --- | --- | --- | --- | --- | --- | --- |
| $T_{\text{inner}}$ | -258040.4949 | 0 | 1 | 1 | 1 | 1 | 1 | 1 | 1 |
| $T_{\text{basal}}$ | -258129.4276 | 88.933 | **0** | **0** | **0** | **0** | **0** | **6.5.10<sup>-15</sup>** | **1.21.10<sup>-71</sup>** |

$\Delta L$ : logLikelihood (logL) difference from the maximal logL in the set.
bp-RELL: bootstrap proportion using REll method (Kishino et al. 1990).
$p$ -KH: p-value of one sided Kishino-Hasegawa test (1989).
$p$ -SH: p-value of Shimodaira-Hasegawa test (2000).
$p$ -WKH: p-value of weighted KH test.
$p$ -WSH: p-value of weighted SH test.
c-ELW: Expected Likelihood Weight (Strimmer & Rambaut 2002).
$p$ -AU: p-value of approximately unbiased (AU) test (Shimodaira, 2002).
All tests performed 10001 resamplings using the REll method. Tests for which the topology was significantly rejected are indicated in bold.

Altogether, our results are therefore congruent with the genomic-based findings recently reported by Bateman et al. (2018) and Piñeiro Fernandez et al. (2019) and contrast with several previous studies. Although being located in a single molecule, plastid regions have been shown to not necessarily behave as a single locus and experience certain forms of intra- and inter-molecular recombination (see Gonçalves et al., 2019; Walker et al., 2019). As most of the plastid genes are also known to encode important biological functions, they may display a sequence evolution patterns that deviate from the species tree topology, and even from non-coding plastid sequences, because of positive selection. Phenomena such as Incomplete Lineage Sorting (ILS), hybridization and introgression, gene duplication of loss as well as horizontal transfers may also affect gene tree topology. Finally, plastids are generally assumed to be maternally inherited in angiosperms but evidences of biparental inheritance have been documented in angiosperms. In spite of all these possible biases potentially affecting plastid gene trees in angiosperms, we did not find any gene strongly supporting the basal position of the *O. insectifera* group in *Ophrys*. We found that particular structural variation also supports the relative phylogenetic proximity of the *O. insectifera* lineage (here *O. aymoninii*) with the *O. sphegodes* relatives (*O. sphegodes* and *O. aveyronensis*). However, *Ophrys aymoninii* shows singular characteristics that further confirms that the *O. insectifera* forms a clearly distinct *Ophrys* lineage. As suggested by other authors for angiosperms, GLS show that the phylogenetic signal of genes such as *matK* slightly outperform *rbcL* but that other *loci* such as *ycf1* (but also *ycf2*), *rpoC2* (see Walker et al., 2019), *rpoB* and *rpoC1* may be considered as good candidates for plastid-based phylogenetic analyses in *Ophrys*. Although our plastid-based findings are congruent with published genome-scale nuclear ones (RADseq- and transcriptome-based) they confirm the necessity of relying on an important number of *loci* to properly infer the evolutionary history of such rapidly evolving groups.

## Acknowledgements

This study is set within the framework of the “Laboratoires d’Excellences (LABEX)” TULIP [ANR-10-LABX-41] and was supported by a ‘Bonus Qualité Recherche’ grant from the Université de Perpignan Via Domitia. We thank Marie-Christine Carpentier, Moaine El Baidouri, Panpan Zhang and the Mechanisms of AdaptatioN and GenOmics (MANGO) team for the support they provided with bioinformatic analyses.

