## Appendix for "Whole plastid genome-based phylogenomics supports an inner placement of the *O. insectifera* group rather than a basal position in the rapidly diversifying *Ophrys* genus (Orchidaceae)"

**Appendix 1 Materials and Methods supplementary information**

**DNA extraction, purification and quantification**

Tubes containing samples were placed in liquid nitrogen and frozen leaves were ground to a fine powder by bead beating on a Silamat S6 (Ivoclar Vivodent, Ellwagen, Germany). The powder was resuspended in 10 mL of 65°C preheated CTAB2X extraction buffer (2% CTAB, 100 mM Tris-HCl pH 8, 20 nM EDTA pH 8, 1.4 M NaCl, 5% N-lauroylsarcosine di-sodium salt, 0.2% 2-mercaptoethanol) and incubated for an hour at 65°C. Then, an equal volume of chloroform was added, and the emulsion was maintained during 10 min before centrifugation at 13,000 rounds per minute (rpm) for 10 min at room temperature. Nucleic acids were precipitated with isopropanol (0.7 v/v) at -80°C for 15 min, centrifugated at 4°C at 13,000 rpm for 10 min. Nucleic acids were further washed with 75% ethanol and centrifugated at 4°C at 13,000 rpm for 10 min. Finally, the pellet was air dried and DNA was resuspended in 300 μL TE (Tris HCl 10 Mm pH 8, EDTA 1 mM pH8) and treated with 100 ng of RNase A (Qiagen, Hilden, Germany) for 30 min at 37°C.

To purify DNA, two steps were added. First, a Genomic DNA Clean and Concentrator-10 column kit (Zymo, Irvine, USA) was used on 5 to 10 μg of DNA treated with RNase A. In a second time, a precipitation with 1/10 volume of sodium acetate 3M pH 5.2 + 2.5 volume of ethanol 100% was performed. The pellet was dissolved in 30 μL of TE and cooled to 4°C overnight. Nanodrop and Qubit (ThermoFisher Scientific, Waltham, USA) quantifications were made to control that concentrations values and reports (260/280 and 260/230 nm ratios) were suitable for MinION (Oxford Nanopore Technologies, Oxford, UK) library preparation and check that the results of the two different quantification methods were similar.

**MinION library preparation and sequencing**

For the two first libraries (one per species), DNA was end-repaired and dA-tailed using NEBNext Ultra II end-repair/dA-tailing module (Biolabs, New England, USA) according to the manufacturer’s instructions except for the thermal cycler program performed with 30 min at 20°C, followed by 30 min at 65°C and terminated with 5 min at 4°C. DNA was further purified with 1X vol. Agencourt AMPure XP beads (Beckman Coulter, High Wycombe, UK). After two washes with 70% ethanol, beads were air dried and the DNA was eluted with 31 μL of distilled water. A Qubit quantification was performed with 1 μL. A ligation was then performed by adding 50 μL Blunt/TAligase master MIX (Biolabs, New England, USA) and 20 μL of Adapter Mix 1D (AMX AD) to the 30 μL A-tailed library and let at room temperature for 30 min. DNA was further purified with 0.4X vol. Agencourt AMPure XP beads (Beckman Coulter, High Wycombe, UK). After two washes with 140 μL of adapter bead binding (ABB), beads were pelleted and air dried a few seconds and DNA was eluted with 15 μL of elution buffer (ELB). A Qubit quantification was performed on 1 μL.

For the two other libraries (one per species) purified DNA was first repaired with NEBNext FFPE DNA repair mix (NEB, Ipswich, USA) for 30 min at room temperature. In a second time, the end-prep step was performed using the NEBNext End-repair/dA-tailing module as per the manufacturer’s instructions except for the thermal cycler program performed for 15 min at 20°C, followed by 5 min at 65°C and 10 min at 70°C, terminated by 5 min at 4°C. A ligation was then performed by adding 50 μL Blunt/TAligase master MIX (Biolabs, New England, USA) and 20 μL of Adapter Mix 1D (AMX AD) to the 30 μL A-tailed library and let at room temperature for 30 min. The DNA was diluted with 100 μL of AMPure Dilution Buffer and transferred to a clean 1,5 mL Eppendorf DNA LoBind tube. It was further purified with 0.25X vol. Agencourt AMPure XP beads during 10 min. After two washes with 250 μL of 0.6x salty adapter bead binding (ABB), and one with 300 μL of TE buffer, beads were pelleted and air dried a few seconds and the DNA was eluted with 14 μL of elution buffer (ELB) and 3 μL of nuclease free water. A Qubit quantification was performed on 1 μL.

MinKNOW application was first used to make the platform QC before starting the analyses and a map/NC-48-h sequencing run flow (Map103) protocol was chosen and initiated. 500 μL of a priming mix (26.6 μL of Fuel Mix (FMX), 500 μL of Running Buffer (RNB), 473.4 μL were introduced twice in the MinION flow cell to prime the sensory array. Thirty six ng of the prepared library was diluted into a mix composed of 75 μL of RNB, 4 μL of FMX and nuclease-free water was added to reach a final volume 150 μL. The 150 μL priming mix was loaded into the sample loading port of the flow cell. After 24 h, 150 μL of a priming mix with 36 ng of the library was loaded again.

**Raw data summary for both short- (Illumina) and -long (Oxford Nanopore Technologies) reads**

| Species | Short-read data (Illumina) | | | Long-read data (Oxford Nanopore Technologies) | | | |
| --- | --- | --- | --- | --- | --- | --- | --- |
|  | Number of reads | Amount of data (G) | Run Accession | Number of reads | Mean read length (bp) | Mean read quality | Run Accession |
| *Ophrys aveyronensis* | 319,740,401 | 95.9 | ERR5101248 | 2,328,146/670,919 | 3,101.8/2,777.1 | 8.9/8.1 | ERR5165940/ ERR5165971/  ERR5166024 |
| *Ophrys aymoninii* | 475,551,312 | 142.7 | ERR5100914 | 2,829,757/1,452,159 | 1,417.6/2,927.8 | 8.0/8.8 | ERR5166026/ ERR5166539 |
| *Ophrys lutea* | 610,990,367 | 183.3 | ERR5303530 | 788,636/118,815 | 5,629.8/10,218.8 | 8.4/8.3 | ERR5167480/  ERR5167484 |
| Raw reads available from the European Nucleotide Archive (ENA). Study Primary Accession PRJEB42431/Secondary Accession ERP12689. Samples: *O. aveyronensis* (Primary Accession ERS5548133/Secondary Accession SAMEA7800875), *O. aymoninii* (ERS5548134/SAMEA7800876) and *O. lutea* (ERS5548135/SAMEA7800877). | | | | | | | |
| For ONT MinION data, two flowcells per sample were used. For *O. aveyronensis* the reads of one of the two flowcells had to be split in two different accessions. | | | | | | | |

**Appendix 2 Characterization of the plastid genomes of *Ophrys aymoninii* and *O. lutea***

| Gene Category | Group of gene | Name of gene |
| --- | --- | --- |
| Self-replication | Ribosomal RNA genes | *rrn16*^a^, *rrn23*^a^, *rrn4.5*^a^, *rrn5*^a^ |
|  | Transfer RNA genes | *trn*A-UGC^a^, *trn*C-GCA, *trn*D-GUC, *trn*E-UUC ^(a)^, *trn*F-GAA, *trn*G-GCC, *trn*H-GUG^a^, *trn*K-UUU, *trn*L-CAA^a^, trnL-UAA, *trn*L-UAG, *trn*M-CAU^b^, *trn*N-GUU^a^, *trn*P-UGG, *trn*Q-UGG, *trn*R-ACG^a^, *trn*R-UCU, *trn*S-CGA, *trn*S-GCU, *trn*S-GGA, *trn*S-UGA, *trn*T-GGU, trnT-UGU, *trn*V-GAC^a^, *trn*V-UAC, *trn*W-CCA, *trn*Y-GUA^(a)^ |
|  | Small subunit of ribosome | *rps*2, *rps*3, *rps*4, *rps*7^a^, *rps*8, *rps*11, *rps*12^a^, *rps*14, *rps*15, *rps*16^c^, *rps*18, *rps*19^a^ |
|  | Large subunit of ribosome | *rpl*2^a^, *rpl*14, *rpl*16, *rpl*20, *rpl*22, *rpl*23^a^, *rpl*32, *rpl*33, *rpl*36 |
|  | DNA-dependent RNA polymerase | *rpo*A, *rpo*B, *rpo*C1^c^, *rpo*C2 |
| Photosynthesis | Subunit of photosystem I (PSI) | *psa*A, *psa*B, *psa*C, *psa*I, *psa*J, *ycf*3^c^, *ycf*4 |
|  | Subunit of photosystem II (PSII) | *psb*A, *psb*B, *psb*C, *psb*D, *psb*E, *psb*F, *psb*H, *psb*I, *psb*J, *psb*K, *psb*L, *psb*M, *psb*N, *psb*T, *psb*Z. |
|  | Subunits of cytochrome b_6_f | *pet*A, *pet*B, *pet*D, *pet*G, *pet*L, *pet*N |
|  | Subunits of ATP synthase | *atp*A, *atp*B, *atp*E, *atp*F^c^, *atp*H, *atp*I |
|  | Subunits of NADH dehydrogenase | *ndh*A^c^, *ndh*B^a,c^, *ndh*C, *ndh*D, *ndh*E, *ndh*F, *ndh*G, *ndh*H, *ndh*I, *ndh*J, *ndh*K |
|  | Large subunits of Rubisco | *rbc*L |
| Other genes | Maturase | *mat*K |
|  | Envelope membrane protein | *cem*A |
|  | Subunit of acetyl-CoA carboxylase | *acc*D |
|  | C-type cytochrome synthesis gene | *ccs*A |
|  | Protease | *clp*P^c^ |
|  | Component of TIC complex | *ycf*1 |
|  | Translation initiation factor IF-1 | *inf*A |
| Genes of unknown function |  | *ycf*2^a^ |

^a^ Duplicated (or possibly duplicated) gene (present in the IR regions)

^b^ Triplicated (or possibly triplicated) gene

^c^ Coding gene containing intron(s)

**Appendix 3** Differences in Gene-wise Log-likelihood scores (ΔGLS) based on genes and CDS

|  | Genes | | | CDS | | |
| --- | --- | --- | --- | --- | --- | --- |
| *Locus* | *T*_inner_ | *T*_basal_ | 𝛥GLS | *T*_inner_ | *T*_basal_ | 𝛥GLS |
| *psbA* | -1546.47 | -1549.28 | 2,81 | -1546.6 | -1549.4 | 2.8 |
| *matK* | -2402.17 | -2406.39 | 4.22 | -2402.34 | -2406.62 | 4.28 |
| *rps16* | -1909.9 | -1920.1 | 10.2 | -389.589 | -389.665 | 0.076 |
| *psbK* | -300.541 | -300.059 | -0.482 | -300.536 | -300.09 | -0.446 |
| *psbI* | -153.904 | -153.905 | 0.001 | -153.902 | -153.903 | 0.001 |
| *atpA* | -2221.17 | -2224.14 | 2.97 | -2221.09 | -2224.04 | 2.95 |
| *atpF* | -2220.46 | -2234.52 | 14.06 | -789.799 | -794.007 | 4.208 |
| *atpH* | -364.919 | -364.653 | -0.266 | -364.874 | -364.619 | -0.255 |
| *atpI* | -1047.8 | -1047.75 | -0.05 | -1047.76 | -1047.72 | -0.04 |
| *rps2* | -1035.04 | -1034.81 | -0.23 | -1035.04 | -1034.84 | -0.2 |
| *rpoC2* | -6352.65 | -6359.53 | 6.88 | -6352.95 | -6359.93 | 6.98 |
| *rpoC1* | -4180.61 | -4193.53 | 12.92 | -3023.98 | -3034.45 | 10.47 |
| *rpoB* | -4661.26 | -4659.8 | -1.46 | -4660.92 | -4659.55 | -1.37 |
| *petN* | -116.89 | -116.891 | 0.001 | -116.889 | -116.889 | 0 |
| *psbM* | -152.632 | -152.623 | -0.009 | -152.622 | -152.613 | -0.009 |
| *psbD* | -1488.67 | -1488.38 | -0.29 | -1488.6 | -1488.33 | -0.27 |
| *psbC* | -2053.75 | -2057.4 | 3.65 | -2053.69 | -2057.3 | 3.61 |
| *psbZ* | -264.976 | -264.74 | -0.236 | -264.988 | -264.764 | -0.224 |
| *rps14* | -481.767 | -488.52 | 6.753 | -481.762 | -488.437 | 6.675 |
| *psaB* | -3217.03 | -3221.41 | 4.38 | -3216.94 | -3221.34 | 4.4 |
| *psaA* | -3289.33 | -3294.54 | 5.21 | -3289.44 | -3294.65 | 5.21 |
| *ycf3* | -2961.21 | -2959.56 | -1.65 | -729.428 | -729.509 | 0.081 |
| *rps4* | -929.109 | -928.359 | -0.75 | -929.01 | -928.304 | -0.706 |
| *atpE* | -581.186 | -581.166 | -0.02 | -581.18 | -581.164 | -0.016 |
| *atpB* | -2166.6 | -2166.56 | -0.04 | -2166.57 | -2166.55 | -0.02 |
| *rbcL* | -2132.08 | -2133.45 | 1.37 | -2132.2 | -2133.39 | 1.19 |
| *accD* | -2241.16 | -2239.78 | -1.38 | -2241.32 | -2240.01 | -1.31 |
| *psaI* | -157.817 | -157.799 | -0.018 | -157.796 | -157.78 | -0.016 |
| *ycf4* | -846.482 | -849.45 | 2.968 | -846.426 | -849.39 | 2.964 |
| *cemA* | -1044.77 | -1047.01 | 2.24 | -1044.69 | -1046.91 | 2.22 |
| *petA* | -1403.16 | -1402.41 | -0.75 | -1403.12 | -1402.4 | -0.72 |
| *psbJ* | -177.744 | -177.735 | -0.009 | -177.734 | -177.725 | -0.009 |
| *psbL* | -157.465 | -157.456 | -0.009 | -157.455 | -157.447 | -0.008 |
| *psbF* | -166.381 | -166.381 | 0 | -166.379 | -166.38 | 0.001 |
| *psbE* | -349.371 | -349.372 | 0.001 | -349.37 | -349.37 | 0 |
| *petL* | -125.975 | -125.976 | 0.001 | -125.974 | -125.974 | 0 |
| *petG* | -166.928 | -166.948 | 0.02 | -166.967 | -166.987 | 0.02 |
| *psaJ* | -252.48 | -252.236 | -0.244 | -252.463 | -252.242 | -0.221 |
| *rpl33* | -276.814 | -276.796 | -0.018 | -276.793 | -276.777 | -0.016 |
| *rps18* | -449.877 | -449.606 | -0.271 | -449.847 | -449.59 | -0.257 |
| *rpl20* | -543.937 | -547.75 | 3.813 | -543.879 | -547.655 | 3.776 |
| *rps12* | -1288.38 | -1288.1 | -0.28 | -530.969 | -530.953 | -0.016 |
| *clpP* | -3741.88 | -3741.82 | -0.06 | -913.901 | -913.623 | -0.278 |
| *psbB* | -2253.92 | -2252.87 | -1.05 | -2253.88 | -2252.9 | -0.98 |
| *psbT* | -174.554 | -173.812 | -0.742 | -174.491 | -173.779 | -0.712 |
| *psbN* | -192.875 | -192.882 | 0.007 | -192.901 | -192.91 | 0.009 |
| *psbH* | -327.917 | -327.89 | -0.027 | -327.886 | -327.861 | -0.025 |
| *petB* | -949.791 | -947.699 | -2.092 | -949.796 | -947.587 | -2.209 |
| *petD* | -719.647 | -719.601 | -0.046 | -719.611 | -719.57 | -0.041 |
| *rpoA* | -1471.77 | -1471.37 | -0.4 | -1471.6 | -1471.24 | -0.36 |
| *rps11* | -650.477 | -650.476 | -0.001 | -650.597 | -650.607 | 0.01 |
| *rpl36* | -167.812 | -167.859 | 0.047 | -167.873 | -167.93 | 0.057 |
| *infA* | -334.821 | -334.794 | -0.027 | -334.79 | -334.765 | -0.025 |
| *rps8* | -557.769 | -557.482 | -0.287 | -557.704 | -557.429 | -0.275 |
| *rpl14* | -567.857 | -573.141 | 5.284 | -567.898 | -573.102 | 5.204 |
| *rpl16* | -641.098 | -640.99 | -0.108 | -640.973 | -640.871 | -0.102 |
| *rps3* | -947.953 | -947.665 | -0.288 | -947.928 | -947.671 | -0.257 |
| *rpl22* | -542.382 | -542.365 | -0.017 | -542.378 | -542.364 | -0.014 |
| *rps19* | -384.164 | -384.156 | -0.008 | -384.154 | -384.146 | -0.008 |
| *rpl2* | -2058.54 | -2058.5 | -0.04 | -1140.65 | -1140.63 | -0.02 |
| *rpl23* | -380.626 | -380.626 | 0 | -380.624 | -380.625 | 0.001 |
| *ycf2* | -10042.9 | -10041.4 | -1.5 | -10042.6 | -10041.3 | -1.3 |
| *rps7* | -636.661 | -636.662 | 0.001 | -636.659 | -636.66 | 0.001 |
| *rpl32* | -417.696 | -416.948 | -0.748 | -416.834 | -416.128 | -0.706 |
| *cssA* | -1408.96 | -1411.78 | 2.82 | -1409.09 | -1411.88 | 2.79 |
| *psaC* | -360.33 | -360.072 | -0.258 | -360.295 | -360.047 | -0.248 |
| *rps15* | -399.624 | -403.341 | 3.717 | -399.595 | -403.283 | 3.688 |
| *ycf1* | -9101.46 | -9121.23 | 19.77 | -9102.18 | -9121.42 | 19.24 |

**Appendix 4** Total length (in bp) of the different CDS of *ndh* subunit genes. Values in bold indicate genes that are consistent with truncation of the associated protein.

|  | *Ophrys iricolor* | *Ophrys lutea* | *Ophrys aymoninii* | *Ophrys sphegodes* | *Ophrys aveyronensis* |
| --- | --- | --- | --- | --- | --- |
| *ndh*A*** | 1092 | 1095 | **66** | 1092 | 1095 |
| *ndhB* | 1533 | 1533 | **120** | **1082** | **1082** |
| *ndhC* | 363 | 363 | **177** | **177** | **177** |
| *ndhD* | 1518 | 1509 | **426** | **588** | **588** |
| *ndhE* | 306 | 306 | **189** | 306 | 306 |
| *ndhF* | **1794** | **220** | *NA* | **293** | **258** |
| *ndhG* | **375** | 531 | **336** | **159** | **159** |
| *ndhH* | 1167 | 1182 | **186** | **585** | **585** |
| *ndhI** | 506 | 486 | **252** | **384** | **369** |
| *ndhJ* | 477 | 477 | **192** | **207** | **207** |
| *ndh*K | 678 | 678 | 678 | **147** | **147** |
| *slight differences in length may be due to differences in annotation between the studies of Roma et al. (2018); Bertrand et al. (2019), and this study. | | | | | |
